# Fyn is Involved in Erythropoietin Signaling Pathway and Interfaces Oxidation to Regulate Erythropoiesis

**DOI:** 10.1101/323873

**Authors:** E Beneduce, A Matte, L De Falco, TSC Mbiandjeu, D Chiabrando, E Tolosano, E Federti, S Petrillo, N Mohandas, A Siciliano, AW Babu, V Menon, S Ghaffari, A Iolascon, L De Franceschi

## Abstract

Erythropoiesis is a complex multistep process responsible of the production of circulating mature erythrocytes and involved the production of reactive oxygen species (ROS) during erythroid differentiation. Here, we document that Fyn, a Src-family-kinase, participates in erythropoietin (EPO) signaling pathway, by the reducing extent of Tyr-phosphorylation of EPO-R and by decreasing STAT5 activity. The importance of Fyn in EPO cascade is also supported by the increased sensitivity of Fyn^−/−^ mice to stress erythropoiesis. Fyn ^−/−^ mouse erythroblasts adapt to the induced stress by the activation of the redox-related-transcription-factor Nrf2. However, the absence of the Nrf2 physiologic repressor Fyn resulted in the persistent activation of Nrf2 and accumulation of non-functional proteins. This is paralleled by ROS induced over-activation of Jak2-Akt-mTOR pathway and repression of autophagy and perturbation of lysosomal-clearance during Fyn ^−/−^ reticulocyte maturation. Treatment with Rapamycin, a mTOR inhibitor and autophagy activator, ameliorates Fyn^−/−^ mouse baseline erythropoiesis and restored the erythropoietic response to phenylhydrazine. Taken together these findings have enabled to identify the novel multimodal action of Fyn in the developmental program of erythropoiesis.

## INTRODUCTION

Erythropoiesis is a complex multistep process during which committed erythroid progenitors undergo terminal differentiation to produce circulating mature red cells. Erythroid differentiation is characterized by the production of reactive oxygen species (ROS) in response to erythropoietin (EPO) and by the large amount of iron imported into the cells for heme biosynthesis.^1^ During erythropoiesis, ROS could function as second messenger by modulating intracellular signaling pathways. EPO induced erythropoiesis activates a signaling cascade, involving Jak2, as the primary kinase, and Lyn, a Tyr-kinase of the Src family (SFK), as secondary kinase.^2–4^ These two kinases target STAT5 transcription factor, one of the key master transcription regulator involved in erythroid maturation events.^2–4^

Previous studies have shown that the mice genetically lacking Lyn (Lyn^−/−^) display reduced STAT5 activation and defective response to phenylhydrazine (PHZ) induced stress erythropoiesis.^2–4^ These data support an important role for Lyn mediated downstream key signaling pathway in EPO induced erythropoiesis.^2–5^ Fyn, is another member of the SFKs that is also expressed in hematopoietic cells.^6–10^ Fyn has been invoked as an additional kinase to the canonical thrombopoietin/Jak2 pathway in megakaryopoiesis.^11^ In addition, Fyn has been shown to target STAT5 and to participate to STAT5 activation in mast-cells in response to FCRI engagement.^8^ Furthermore, Fyn intersects different intracellular signaling pathways and participates to the regulation of the redox sensitive transcriptional factor Nrf2.^12–14^ Nrf2 regulates the expression of a number of genes involved in cellular antioxidant and cytoprotective systems and thereby enables the cell to protect against oxidation damage.^12–14^ Following acute phase response, Fyn switches-off active Nrf2, triggering its exit from the nucleus and degradation.^12–15^ In erythroid maturation events, the activation of Nrf2 is crucial to support stress erythropoiesis induced by the oxidant, PHZ, and in modulating ineffective erythropoiesis in β-thalassemic mice.^16,17^ In other cellular models, it has been shown that impairment of Nrf2 post-induction regulation results in perturbation of cell homeostasis and in accumulation of poly-ubiquitylated protein aggregates due to deregulated autophagy.^14^ Autophagy is activated in response to different cellular stresses to ensure cell survival and ensure the clearance of the damaged proteins.^16,17^ We recently showed that in chorea-acanthocytosis the impairment of autophagy promotes accumulation of proteins, resulting in engulfment of the cells and in perturbation of erythropoiesis combined with increased oxidative stress.^18^ Thus, the response to EPO-R activation might involve a integrated and intersecting actions of a number of intracellular signaling pathways The role of these intersecting pathways might include modulation of Nrf2 function to control ROS generation and to clear damage proteins by autophagy.

In present study, we explored the role of Fyn in regulating normal and stress erythropoiesis. We show that in addition to Jak2 and Lyn, Fyn is an additional kinase involved in EPO signaling cascade by targeting STAT5 activation. The reduction in the efficiency of EPO signal promotes the generation of ROS and the over-activation of Jak2-Akt-mTOR pathway and the repression of autophagy. The absence of Fyn results in persistent activation of Nrf2 and accumulation of damaged proteins. This is further amplified by the blockage of autophagy mediated by mTOR activation, which markedly perturbs the response to stress erythropoiesis induced by either phenylhydrazine (PHZ) or Doxorubicin. In Fyn^−/−^ mice, the rescue experiments with Rapamycin, an mTOR inhibitor and autophagy activator, co-administrated to PHZ further validated the importance of autophagy as adaptive mechanism to stress erythropoiesis in presence of perturbation of EPO cascade.

## METHODS

### Mouse strains and design of the study

The Institutional Animal Experimental Committee of University of Verona (CIRSAL) and the Italian Ministry of Health approved the experimental protocols. Two-months old female wild-type (WT) and Fyn^−/−^ mice were studied. Where indicated, WT and Fyn^−/−^ mice were treated with EPO (10 U/mouse/day for 5 days by intraperitoneal injection),^3^ or Phenylhydrazine (PHZ: 40 mg/Kg on day 0 by intraperitoneal injection),^17^ or Doxorubicin (DOXO: 0.25 mg/Kg on day 0 by intraperitoneal injection)^19^ to study stress erythropoiesis. Rapamycin (Rapa) was administrated at the dosage of 10 mg/Kg/d by intraperitoneal injection for 1 week, then mice were analyzed. In experiments with PHZ co-administration, Rapa was given at the dosage of 10 mg/Kg/d by intraperitoneal injection one day before PHZ administration (40 mg/Kg body; single dose at day 0) and then Rapa was maintained for additional 14 days. N-Acetylcysteine (NAC, 100 mg/Kg body; intraperitoneally injected) was administrated for 3 weeks as antioxidant treatment.^16,17^ In mouse strains, hematological parameters, red cell indices and reticulocyte count were evaluated at baseline and at different time points (6, 8 and 11 days after EPO injection; at 2, 4, 8 and 14 days after PHZ injection; at 3, 6 and 9 days after DOXO injection; at 2, 4, 8, 14 days after Rapa plus PHZ injection) as previously reported.^20,21^ Blood was collected with retro-orbital venipuncture in anesthetized mice using heparinized microcapillary tubes. Hematological parameters were evaluated on a Bayer Technicon Analyser ADVIA. Hematocrit and hemoglobin were manually determined.^22,23^

### Flow cytometric analysis of mouse erythroid precursors and molecular analysis of sorted erythroid cells

Flow cytometric analysis of erythroid precursors from bone marrow and spleen from WT and Fyn^−/−^ was carried out as previously described using the CD44-Ter119 or CD71-Ter119 strategies.^16,24,25^ Analysis of apoptotic basophilic, polychromatic and orthochromatic erythroblasts was carried out on the CD44-Ter119 gated cells using the Annexin-V PE Apoptosis detection kit (eBioscience, San Diego, CA, USA) following the manufacturer’s instructions. Erythroblasts ROS levels were measured as previously reported by Matte et al.^16^ Sorted cells were used for (i) morphological analysis of erythroid precursors on cytospin preparations stained with May Grunwald-Giemsa; (ii) immuno-blot analysis with specific antibodies against anti-P-Ser473-Akt, anti-Akt, anti-P-Ser2448-mTOR, anti-mTOR, anti-Jak2 (Cell Signaling, Massachusetts, USA); anti-P-Ser40-Nrf2, anti-Nrf2, anti-p62, anti-Rab5 (Abcam, Cambridge, UK); anti-Keap1 (Proteintech, Manchester, UK); anti-EPO-R (Sigma-Aldrich, Missouri, USA); anti-STAT5, anti-Lyn (Santa Cruz Biotechnology, Texas, USA); anti-GAPDH (Santa Cruz Biotechnology, Texas, USA) and anti-catalase (Abcam, Cambridge, UK) were used as loading control; (iii) immunoprecipitation assay; and (iv) RT-PCR analysis. Details of immunoprecipitation, RT-PCR and immuno-blot protocols used for the analysis of sorted erythroblasts are described in Supplementary materials and methods.

### CFU-E, BFU-E assay

CFU-E and BFU-E assay was carried out using MethoCult as previously reported.^26^ Details are present in Supplementary Methods.

### Immunofluorescence assay for p62 and FOXO3 in sorted erythroblasts

Immunofluorescence assay for p62 and FOXO3 in sorted erythroblasts was carried out as previously described.^18,23,27^ Details are reported in Supplementary materials and methods.

### LysoTracker and MitoTracker analysis in maturating reticulocytes

To obtain reticulocyte enriched RBC fraction, WT and Fyn^−/−^ mice were intraperitoneally injected with PHZ (40 mg/kg) at day 0, 1, 3 to induce reticulocytosis, and blood was collected in heparinized tubes at day 7, as previously described.^28^ RBCs were washed three times with the maturation medium (60% IMDM, 2mM L-glutamine, 100U Penicillin-Streptomicin, 30% FBS, 1% BSA and 0.5 μg/ml Amphotericin), diluted 1/500 in maturation medium and cultured in a cell culture incubator at 37°C, 5% of CO_2_ for 3 days. Clearance of Lysosome and Mitochondria, on the CD71/Ter119 gated RBC populations, were analyzed at day 0 and 3 of culture using the Lysotracker Green DND-26 (ThermoFisher Scientific) and the MitoTracker Deep Red (ThermoFisher Scientific) probes, respectively, following the manufacturer’s instructions. Samples were acquired using the FACSCantoN™ flow cytometer (Becton Dickinson, San Jose, CA, USA) and data were processed with the FlowJo software (Tree Star, Ashland, OR, USA) as previously described.^16,17^

### Pearl’s analysis of liver and spleen

Immediately following dissection, spleen and liver were formalin-fixed and paraffin-embedded for Pearl’s staining.

### Molecular analysis of liver

Protocols used for RNA isolation, cDNA preparation and quantitative qRT-PCR have been previously described.^29^ Detailed primer sequences are available on request and shown in Table 1S. Liver immuno-blot analysis was performed as previously described.^16,30^

### Measurement of heme and heme-oxygenase-1 activity

Liver heme content was measured using a fluorescence assay, as previously reported.^31^ Details are reported in Supplementary Methods.

HO activity was evaluated in tissue microsomal fractions by spectrophotometric determination of bilirubin produced from hemin added as the substrate, as previously reported.^32^

### Statistical analysis

Data were analyzed using either t-test or the 2-way analysis of variance (ANOVA) for repeated measures between the mice of various genotypes. A difference with a p< 0.05 was considered significant.

## RESULTS

### The absence of Fyn results in decreased efficiency of EPO-signaling pathway

Fyn^−/−^ mice displayed a slight microcytic anemia characterized by a small but significant reduction in hemoglobin, microcytosis and increased reticulocyte counts compared to wild-type animals (Table 1). To understand whether iron deficiency might account for the observed microcytosis, we evaluated iron accumulation in the liver and spleen. No differences in Pearl’s staining for iron in either liver or spleen of Fyn^−/−^ compared to wild type mice was observed (Figure 1Sa). In agreement, expression levels of H-Ferritin in liver were similar in both mouse strains, whereas expression of L-Ferritin was slightly, but significantly lower in Fyn^−/−^ mice compared to wild-type mice (Figure 1Sb). Haptoglobin levels were measured to determine the possible contribution of hemolysis to microcytic anemia in Fyn^−/−^ mouse. Up-regulation of haptoglobin mRNA levels was noted in liver from Fyn^−/−^ mice, while plasma haptoglobin levels were similar in both mouse strains (Figure 1Sc, d). These findings suggest that in mice genetically lacking Fyn, the noted mildly compensated anemia is not related to either iron deficiency or chronic hemolysis.

**Table 1.** Hematological Parameters and Red Cell Indices in Wild-type and Fyn^−/−^ Mice.

|  |  |  |
| --- | --- | --- |
|  | **Wildtype mice**<br>**(n=15)** | **Fyn<sup>-/-</sup> mice**<br>**(n=15)** |
| Hct (%) | 48.2 ± 1.3 | 46.1 ± 0.8* |
| Hb (g/dl) | 15.9 ± 0.6 | 14.3 ± 0.5* |
| MCV (fl) | 50.3 ± 0.4 | 46.5 ± 1.3* |
| MCH (g/dl) | 15.3 ± 0.3 | 14.8 ± 1.1 |
| RDW (%) | 11.6 ± 0.3 | 13.2 ± 0.4* |
| Retics (10<sup>3</sup> cells/uL) | 451 ± 40.7 | 559 ± 45* |

Hct: hematocrit; Hb: hemoglobin; MCV: mean corpuscular volume; MCH: mean corpuscular hemoglobin;
RDW: red cell distribution width; Retics: reticulocytes; \*p< 0.05 compared to wild-type mice;

To better define the Fyn^−/−^ mouse hematologic phenotype, we carried out the morphologic analysis of erythroblasts at distinct stages of terminal erythroid differentiation. As shown in Figure 1a, decreased chromatin condensation and larger cellular size was a characteristic feature of different populations of sorted Fyn^−/−^ mouse erythroblasts (pop II: basophilic erythroblasts; pop III: polychromatic erythroblasts and pop IV: orthochromatic erythroblasts; Figure 1a). Furthermore, an increase in number of total erythroblasts in bone marrow was noted, without evidence of extramedullary erythropoiesis (Figure 1b). The maturation profile of erythroblasts revealed an accumulation of orthochromatic erythroblasts (Figure 2Sa). When CD44/Ter119 approach was used to characterize erythropoiesis, no major differences in either total erythroblasts or in erythroblasts subpopulations between wild-type and Fyn^−/−^ mice were observed (Figure 2Sb, c). Up-regulation of erythropoietin (EPO) gene expression in kidney was found in Fyn^−/−^ mice compared to that of wild-type animals (Figure 2Sd). In addition, we found increased ROS levels throughout Fyn^−/−^ erythroid maturation from basophilic erythroblasts (pop II) to polychromatic (pop III) and orthochromatic erythroblasts (pop IV) compared to wild-type cells (Figure 1c, upper panel). This was associated with higher amounts of Annexin V^+^ cells in the different subpopulation of erythroblasts compared to wild-type cells (Figure 1c], lower panel). Collectively, these findings indicate a decreased efficiency of EPO signaling pathway in the absence of Fyn. To understand the impact of Fyn on EPO cascade, we evaluated the EPO-Jak2-STAT5 signaling pathway in sorted Fyn^−/−^ erythroblasts. As shown in Figure 1d, reduced activation of EPO-Receptor (EPO-R) as reflected by decreased EPO-R Tyr-phosphorylation noted in erythroblasts genetically lacking Fyn (Figure 1d). This was associated with increased activation of Jak2 kinase without any change in Lyn activity compared to wild-type cells (Figure 1d). In agreement with the reduction in EPO-R activation, we observed a significant decrease in STAT5 with concomitant down-regulation of *Cish* expression, a well-documented gene target of STAT5 in sorted Fyn^−/−^ erythroblasts (Figure 1d; Figure 2Se). Following treatment with recombinant EPO (10 U/day for 5 days), Fyn^−/−^ mice showed a blunted response in terms of increases in Hct and reticulocyte counts compared to wild-type animals (Figure 1e).

**Figure 1.**
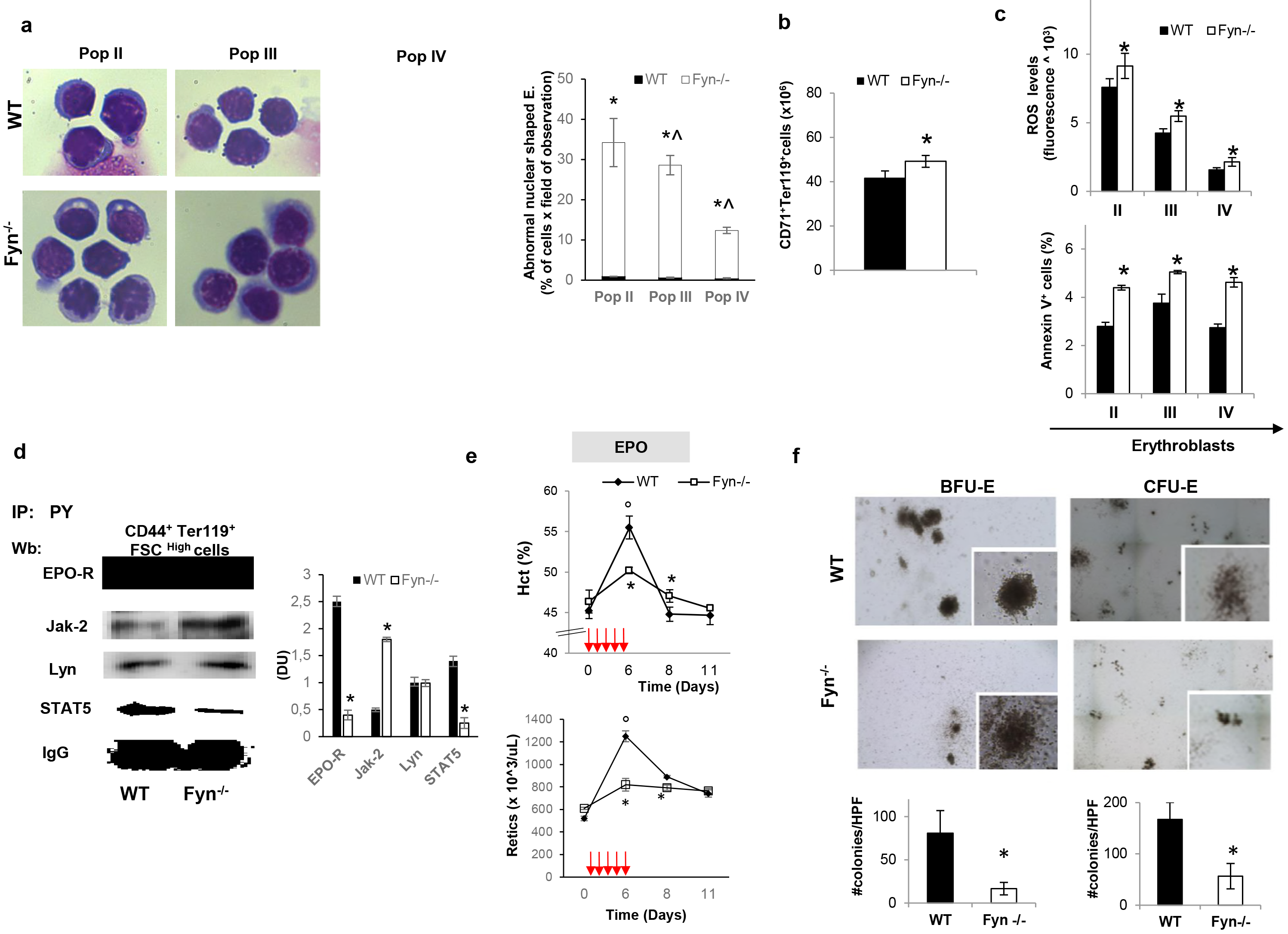
The absence of Fyn results in perturbation of EPO signaling cascade. **(a) Left panel.** Morphology of sorted erythroid precursors: population II (Pop II), corresponding to basophilic erythroblasts; population III (Pop III), corresponding to polychromatic erythroblasts and population IV (Pop IV), corresponding to orthochromatic erythroblasts, from bone marrow of wild-type (WT) and Fyn^−/−^ mice. Cytospins were stained with May-Grunwald-Giemsa. Cells were imaged under oil at 100x magnification using a Panfluor objective with 1.30 numeric aperture on a Nikon Eclipse DS-5M camera and processed with Digital Slide (DS-L1) Nikon. One representative image from a total of 10 for each mouse strains. **Right panel.** Abnormal nuclear shaped erythroblasts and binuclear erythroblasts from WT and Fyn ^−/−^ mice evaluated on cytospin stained with May-Grunwald-Giemsa. Data are presented as means ±SD (*n*=8 from each strain); *p<0.05 compared to WT; ^ p <0.05 compared to Pop II^-^. **(b)** Cyto-fluorimetric analysis of total erythroid precursors from the bone marrow of wild-type (WT) and Fyn^−/−^ mice using the following surface markers: CD71 and Ter119 (see also the Supplementary Materials and Methods and Figure 2Sa for maturation profile). Data are presented as means ±SD (*n*=8); * p < 0.05 compared to WT. **(c) Upper panel.** ROS levels in erythroid precursors: population II (Pop II), corresponding to basophilic erythroblasts; population III (Pop III), corresponding to polychromatic erythroblasts and population IV (Pop IV), corresponding to orthochromatic erythroblasts from bone marrow of wild-type (WT) and Fyn^−/−^ mice. Data are presented as means ±SD (*n*=10 from each strain); * p <0.05 compared to WT. **Lower panel.** Amount of Annexin V^+^ cells in Pop II, III and IV from bone marrow of wild-type (WT) and Fyn^−/−^ mice. Data are presented as means ±SD (*n*=8 from each strain); * p <0.05 compared to WT. **(d)** Total Tyrosin-(Tyr) phosphorylated proteins were immunoprecipitated from bone marrow sorted erythroblasts of wild-type (WT) and Fyn^−/−^ mice and detected with antibody to erythropoietin- receptor (EPO-R), Janus kinase-2 (Jak-2), Lyn kinase (Lyn), Signal transducer and activator of transcription 5 (STAT5). The experiment shown is representative of 6 such experiments. IgG was used as loading control. **Right panel.** Densitometric analyses of the immunoblot bands similar to those shown are presented at right (DU: densitometric Unit). Data are shown as means ±SD (*n*=6; *p<0.01 compared to WT). **(e)** Hematocrit (%) and reticulocyte count in wild-type (WT; *n*=6) and Fyn^−/−^ (*n*=6) mice exposed to recombinant erythropoietin injection (EPO 50 U/Kg/die, red arrows). Data are presented as means ±SD; *p< 0.05 compared to WT mice; °p<0.05 compared to baseline values. **(f)** To assess the number of erythroid progenitor bone marrow derived cells were cultured in vitro under erythroid osteogenic differentiation conditions. The CFU-E and BFU-E from wild-type (WT) and Fyn^−/−^ mice were quantified (#CFU-E or BFU-E/dish; **lower panel)**; data are shown as means ±SD (n=6; p <0.05 compared to WT).

To explore whether the reduced efficiency of EPO cascade might also involve erythroid progenitors, we carried out the *in vitro* erythroid cell colony forming assay. A lower number of CFU-E and BFU-E colony forming cells were found in Fyn^−/−^ bone marrow (Figure 1f). This was associated again with lower activation of EPO-R mediated signaling cascade with a reduced activation of STAT5 but hyper-activation of Jak2 in Fyn^−/−^ CFU-E (Figure 2Sf).

Our data indicate that Fyn is involved in EPO signaling cascade and the increased ROS generation contributes to the hyper-activation of Jak2 in presence of reduced efficiency of EPO signaling pathway.^33^ Thus, the very mild microcytic anemia phenotype of Fyn ^−/−^ mice is likely to be related to reduced STAT5 activation, as observed in mice genetically lacking STAT5 than to perturbation of iron metabolism.^34^

### Fyn^−/−^ mice display increased sensitivity to PHZ or doxorubicin induced stress erythropoiesis

Since EPO is the primary signal in stress erythropoiesis, we treated Fyn^−/−^ mice with either PHZ to induce acute hemolytic anemia due to severe oxidative stress or Doxorubicin that temporary represses erythropoiesis with generation of ROS.^17,19^ PHZ treatment induced a similar drop in Hct levels in both mouse strains at day 2 following PHZ administration (Figure 2a, upper panel). However, the Hct and reticulocyte recovery were blunted in Fyn^−/−^ mice compared to control animals (Figure 2a, upper and lower panel). Extramedullary erythropoiesis as assessed by increased splenic erythropoiesis showed a blunted response in Fyn^−/−^ mice at day 4 following PHZ treatment with a compensatory increase by day 14 (Figure 2b, upper panel, see also Figure 3Sa for absolute values of number of erythroblasts at day 4 after PHZ). In bone marrow, we observed a mild increase in total erythroblasts in both mouse strains at day 2 and 4 after PHZ injection (Figure 2b, lower panel). It is of interest to note that in Fyn^−/−^ mice at day 8 following PHZ treatment, we observed a significant increase in the total number of bone marrow erythroblasts as possible compensatory mechanism due to the failure in efficient activation of splenic extramedullary erythropoiesis (Figure 2b lower panel). The amount of Annexin V^+^ cells following PHZ treatment was higher in Fyn^−/−^ polychromatic and orthochromatic erythroblasts compared to wild-type cells (Figure 2c).

**Figure 2.**
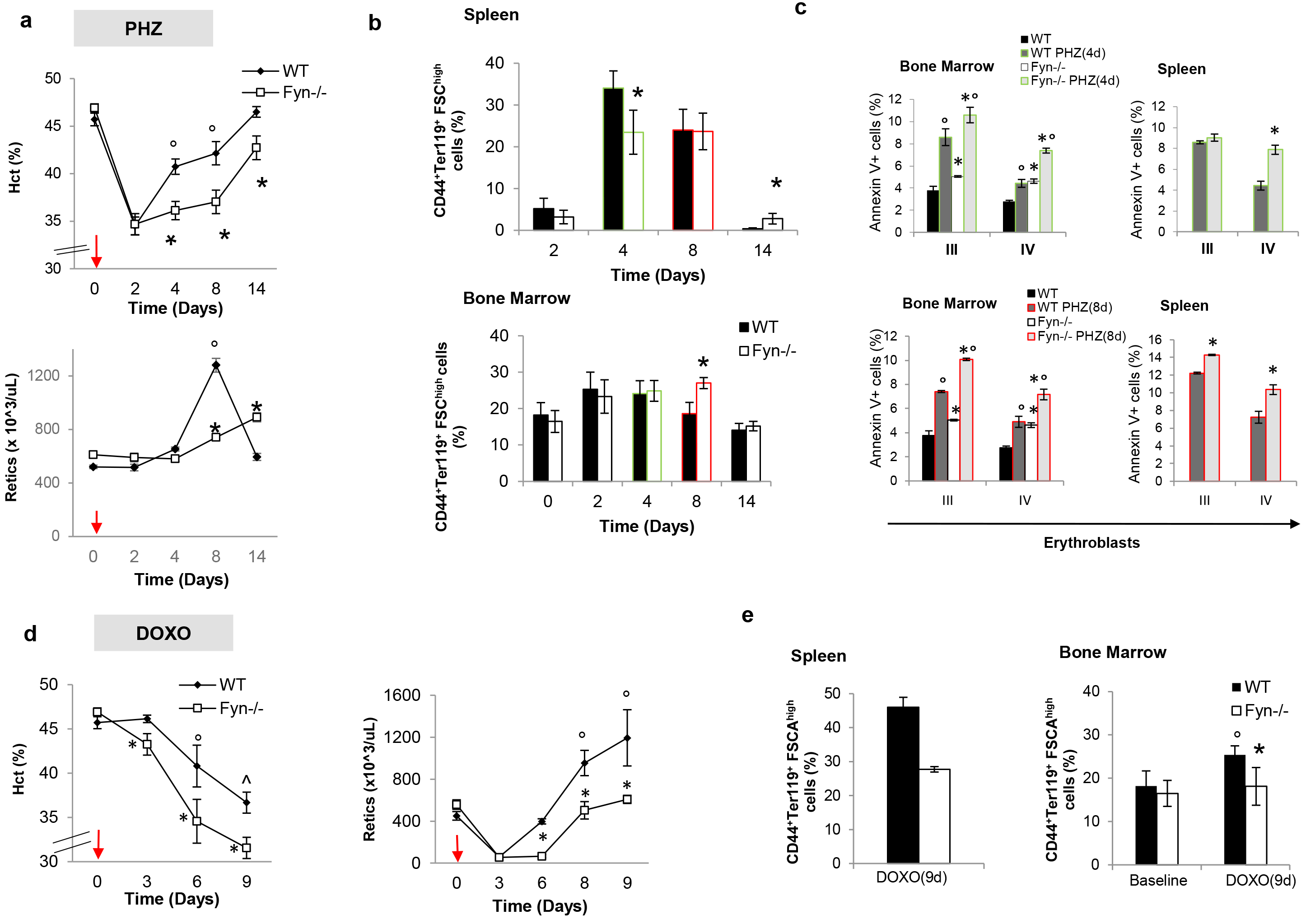
A blunted response to stress erythropoiesis characterizes Fyn^−/−^ mice. **(a)** Hematocrit (%) and reticulocyte count in wild-type (WT; *n*=6) and Fyn^−/−^ (*n*=6) mice exposed to phenylhydrazine injection (PHZ /Kg/die, red arrows). Data are presented as means ±SD; *p< 0.05 compared to WT mice; °p<0.05 compared to baseline values. **(b)** Cyto-fluorimetric analysis of total erythroid precursors from the bone marrow and the spleen of wild-type (WT) and Fyn^−/−^ mice using the following surface markers: CD44 and Ter119 (see also the Supplementary Materials and Methods and Figure 3Sa for absolute values). Data are presented as means ±SD (*n*=6); * p < 0.05 compared to WT. Since we focus on day 4 and day 8 after PHZ administration, we highlighted them respectively in green and red. This color code is used also in Figure 2c. **(c)** Amount of Annexin-V^+^ cells in population III (Pop III), corresponding to polychromatic erythroblasts and population IV (Pop IV), corresponding to orthochromatic erythroblasts from either spleen or bone marrow of wild-type (WT) and Fyn^−/−^ mice respectively at 4 (green) and 8 (red) days after PHZ administration. Data are presented as means ±SD (*n*=6 from each strain); \**P*<0.05 compared to WT. **(d)** Hematocrit (%) and reticulocyte count in wild-type (WT; *n*=6) and Fyn^−/−^ (*n*=6) mice exposed to doxorubicin injection (DOXO 0,25 mg/Kg/die, red arrows). Data are presented as means ±SD; *p< 0.05 compared to WT mice; °p<0.05 compared to baseline values. **(e)** Cyto-fluorimetric analysis of total erythroid precursors from the bone marrow and the spleen of wild-type (WT) and Fyn^−/−^ mice using the following surface markers: CD44 and Ter119 (see also the Supplementary Materials and Methods and Figure 3Sb for absolute values) 9 days after Doxorubicin injection. Data are presented as means ± SD (*n*=6); * p < 0.05 compared to WT; °p<0.05 compared to baseline values.

Doxorubicin induced a more severe and prolonged anemia in Fyn^−/−^ mice than in wild-type animals (Figure 2d, left panel). At day 3 and 6 following Doxorubicin treatment, we noted a plateau in reticulocyte count in Fyn^−/−^ mice (Figure 2d]), suggesting a substantial impairment in the reticulocyte response compared to Doxorubicin treated wild-type animals. Enumeration of total number of erythroblasts in spleen and bone marrow at day 9 after Doxorubicin administration, showed a substantial reduction in both bone marrow and splenic erythropoiesis in Fyn^−/−^ mice compared to wild-type animals (Figure 2e; see also Figure 3Sb for absolute values). Increases in the numbers of Annexin V^+^ polychromatic and orthochromatic erythroblasts was noted in Fyn^−/−^ mice compared to wild-type animals at 9 days after Doxorubicin administration (Figure 3Sc). The findings of increased sensitivity of Fyn^−/−^ mice to stress erythropoiesis induced by PHZ or Doxorubicin, further validate the importance of Fyn in EPO signaling cascade.

### Increased activation of Akt in Fyn^−/−^ mice contributes to the modulation of redox cellular response during erythropoiesis

In normal and disordered erythropoiesis, previous studies have shown that Jak2 and oxidation can activate Akt, which affects multiple targets during erythropoiesis (Figure 3a).^4^ Notably, Akt is also important in mediating cellular response to oxidation by the activation of two redox sensitive transcriptional factors, Nrf2 and Forkhead box-O3 (FOXO3), as well as of mTOR, the gatekeeper of autophagy activation.^35–37^ Fyn^−/−^ mouse erythroblasts displayed higher levels of active Akt (Ser 473) compared to wild-type cells (Figure 3b) in association with increased activation of Nrf2, as indicated by higher phospho-Nrf2 form in Fyn^−/−^ mouse erythroblasts compared to wild-type cells (Figure 3c). The activation of FOXO3 was evaluated by both immunomicroscopy and immunoblot analysis, this latter using a specific antibody against inactive phospho-FOXO3. In Fyn^−/−^ mouse erythroblasts, we found a slight but not significant increase in activation of FOXO3 compared to wild-type erythroblasts (Figure 4Sa). We then focused on Nrf2 since Fyn is important in postinduction regulation of Nrf2.^14^ The up-regulation of ARE-genes for anti-oxidant systems such as catalase, Gpx1, HO1 and Prx2 confirmed the increased Nrf2 function in Fyn^−/−^ mouse erythroblasts (Figure 3d). Immunoblot analysis with specific antibodies for the corresponding proteins further validated the activation of Nrf2 pathway (Figure 4Sb).

**Figure 3.**
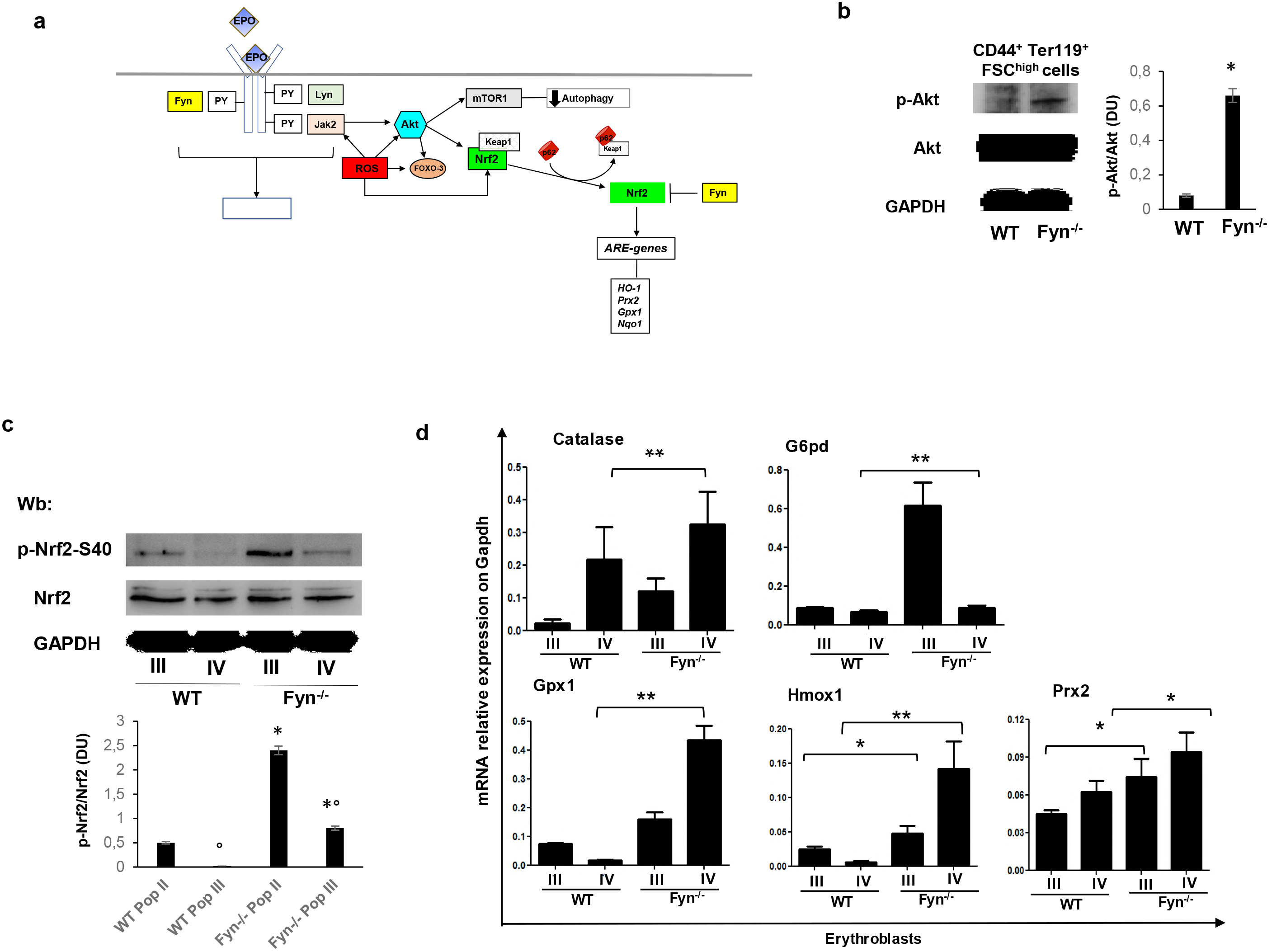
Fyn^−/−^ mice display the activation of Akt and the redox related transcription factor Nrf2. **(a)** Schematic diagram of erythropoietin (EPO) cascade in Fyn^−/−^ mice. Fyn acts as additional kinase to Jak2 and Lyn in EPO signaling pathway. In other models of stress erythropoiesis, increase oxidation contributes to the activation of Jak2-Akt-mTOR system, resulting in repression of autophagy. Akt might activate different redox sensitive transcription factors such as Forkhead box-O3 (FOXO3) or Nrf2. Nrf2 activation requires its release from the shuttle protein Keap1 and its nuclear translocation as phosho-Nrf2. This up-regulates the ARE-genes for cytoprotective systems such as heme-oxygenase-1 (HO-1), peroxiredoxin-2 (Prx2), glutathione peroxidase 1 (Gpx1) or NAD(P)H quinone dehydrogenase 1 (Nqo1). Since Nrf2 post-induction regulation is Fyn-dependent, we hypothesized a dysregulation of Nrf2, possibly contribution to the blunted response of Fyn^−/−^ to stress erythropoiesis. Oxidation and Akt might also contribute to the activation of the redox sensitive transcription factor Forkhead box-O3 (FOXO3). **(b)** Western-blot (Wb) analysis of phospho-Akt (p-Akt) and Akt in sorted CD44^+^Ter119^+^FSC^high^ bone marrow cells from wild-type (WT) and Fyn^−/−^ mice. GAPDH was used as protein loading control. Densitometric analyses of the immunoblot bands similar to those shown are presented at right. Data are shown as means ±SD (*n*=6; *p<0.01 compared to WT). **(c)** Western-blot (Wb) analysis of phospho-Nrf2 (Serine 40, S40) and Nrf2 in sorted erythroblasts: population III (Pop III), corresponding to polychromatic erythroblasts and population IV (Pop IV), corresponding to orthochromatic erythroblasts, from wild-type (WT) and Fyn^−/−^ mice. GAPDH was used as protein loading control. **Lower panel.** Densitometric analyses of the immunoblot bands similar to those shown are presented at right (DU: densitometric Unit). Data are shown as means ±SD (*n*=6; *p<0.01 compared to WT). **(d)** RT-PCR expression of heme-oxygenase-1 (HO-1), peroxiredoxin-2 (Prx2), glutathione peroxidase 1 (Gpx1), glucose-6-phosphate dehydrogenase (G6PD) and catalase on sorted mouse polychromatic (Pop III) and orthochromatic (Pop IV) erythroblasts from bone marrow of wild-type and Fyn^−/−^ mice. Experiments were performed in triplicate. Error bars represent the standard deviations (mean ±SD); \**p*<0.05, \*\**p* <0.001.

However, the increased expression of anti-oxidant and cyto-protective system related to Nrf2 function seems unable to completely counteract induced oxidative damage in Fyn^−/−^ mouse erythroblasts. Overall these effects are similar to those observed in both cell and animal models characterized by prolonged Nrf2 activation, which is associated with severe and even lethal phenotype,^13,38^ mainly related to the accumulation of damaged proteins and the perturbation of autophagy.

Among the many Nrf2 related genes up-regulated in Fyn^−/−^ mouse erythroblasts, we focused on HO-1, the main heme-catabolizing enzyme under stress conditions and a major player in the maintenance of cell homeostasis.^39^

### Heme-oxygenase activity and heme levels are similar in Fyn^−/−^ and wild-type mice

Since systemic heme homeostasis is orchestrated by the liver, we evaluated the impact of Fyn deficiency on hepatic heme catabolism. First, we confirmed the increased activation of Nrf2 in Fyn^−/−^ liver compared to wild-type counterpart (Figure 5Sa). When we analyzed HO-1 expression and HO-1 activity in this organ, despite similar levels of HO-1 mRNA in the livers of wild-type and Fyn^−/−^ mice, HO-1 protein level was higher in the Fyn^−/−^ mice. Similar findings were also noted in erythroid cells (Figure 5Sb). Interestingly, in spite of increased protein levels, hepatic HO activity was unchanged in Fyn^−/−^ animals (Figure 5Sc) suggesting no alteration in heme catabolism. This conclusion was supported by the finding that the heme content in the liver that was similar between wild-type and Fyn^−/−^ mice (Figure 5Sd). Thus, in the absence of Fyn HO-1 protein is increased but its activity is unchanged in Fyn^−/−^ mice, suggesting the accumulation of functionally inactive HO-1.^14^ Autophagy is the master control system regulating protein quality and clearance of damaged proteins.^35,37,40^ The lysosomal related cargo p62 protein can be used as indirect marker of autophagy and its accumulation correlates with impairment of autophagy.^41^ In liver from Fyn^−/−^ mice, we found an accumulation of p62, suggesting a blockage of autophagy in the absence of Fyn (Figure 5Se, f). In agreement, mTOR was more active in liver from Fyn^−/−^ mice compared to wild-type animals (Figure 5Sg).^35,42,43^ These data imply impaired autophagy in liver from mice genetically lacking Fyn.

### Impaired autophagy related to mTOR activation characterizes Fyn^−/−^ mouse erythropoiesis

Since autophagy is also important during erythropoiesis,^40^ we explored mTOR signaling during erythroid maturation in Fyn^−/−^ mouse. As shown in Figure 4a, Fyn^−/−^ mouse erythroblasts displayed increased activation of phosho-mTOR compared to wild-type cells in association with accumulation of p62, similarly to that noted in Fyn^−/−^ mouse liver (Figure 4a) as well as of Rab5, a small GTP protein involved in the late phases of autophagy (Figure 4a).^37^ Consistent with a blockage of autophagy during erythropoiesis in Fyn^−/−^ mouse, we noted accumulation of p62 in large clusters in Fyn^−/−^ erythroblasts compared to wild-type cells (Figure 4b, Figure 6Sa). Since p62 acts as autophagy adaptor, controlling proteins turnover,^13,14^ we evaluated Keap1, a known substrate of p62.^14,41^ In Fyn^−/−^ erythroblasts, we found increased accumulation of Keap1 compared to wild-type cells (Figure 6Sb). Co-immunoprecipitation with either antibodies to p62 (Figure 4c) or Keap1 (Figure 4d) showed accumulation of p62- Keap1 complex in Fyn^−/−^ mouse erythroblasts. If the perturbation of autophagy in Fyn^−/−^ mice is physiologically relevant to erythroid maturation, it is also likely to affect reticulocyte maturation.^28,40,44^ To test this possibility, we evaluated the *in vitro* maturation of reticulocytes from PHZ treated mice.^28^ As shown in Figure 5a, decreased maturation of Fyn^−/−^ mouse reticulocyte was detected as assessed by decreased lysosomal clearance detected by the LysoTracker analysis (Figure 5b). Interesting, no difference in mitochondrial clearance using MitoTracker analysis was noted (Figure 5b, lower panel). Documentation of increased accumulation of p62 in Fyn^−/−^ reticulocyte further supports the impairment of autophagy during erythroid maturation in Fyn^−/−^ mice (Figure 5c).

**Figure 4.**
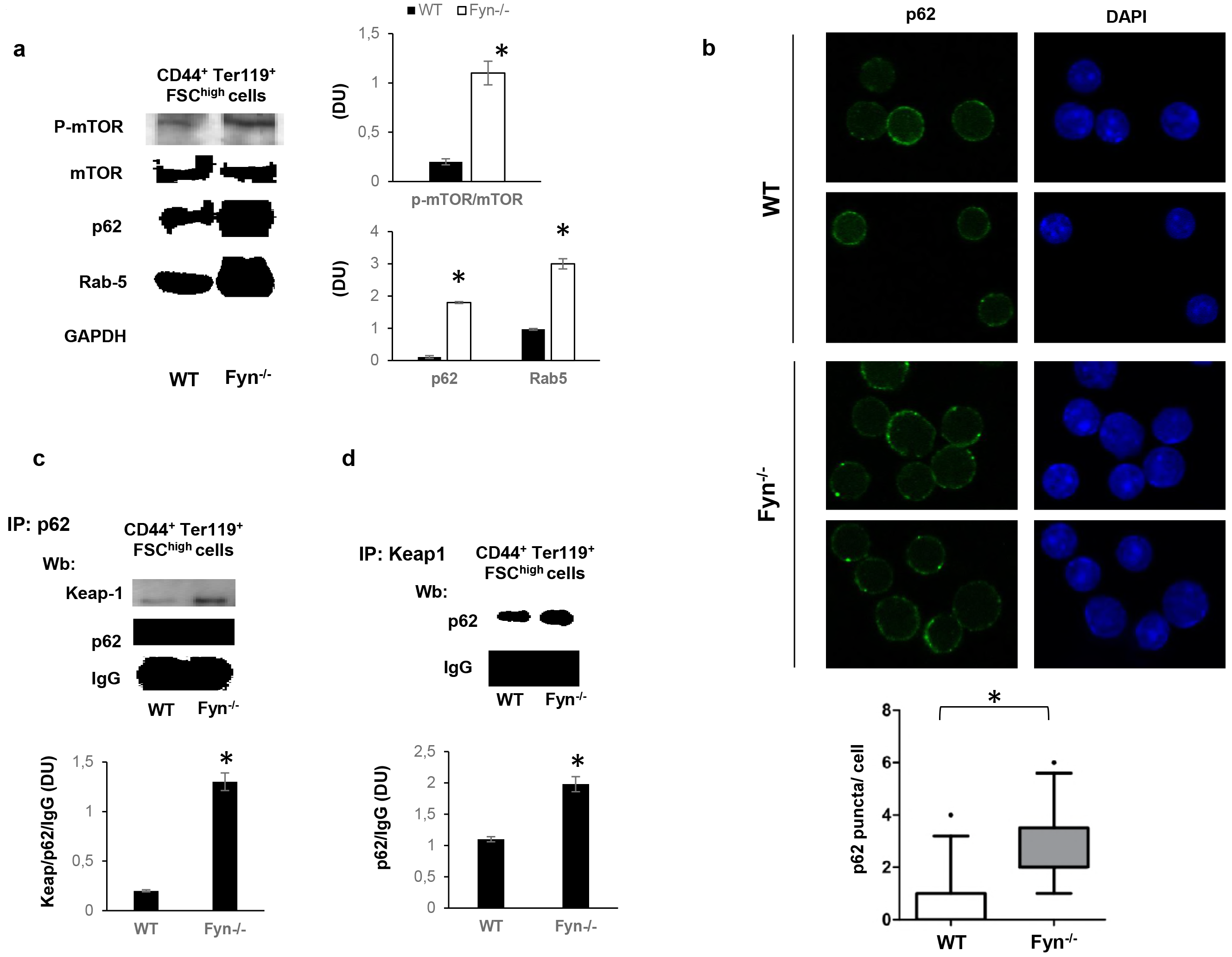
Activation of mTOR and impaired autophagy characterize Fyn^−/−^ mouse erythroblasts. **(a)** Western-blot (Wb) analysis of phospho-mTOR (p-mTOR), m-TOR, p62 and Rab 5 in sorted CD44^+^Ter119^+^FSC^High^ bone marrow cells from wild-type (WT; *n*=4) and Fyn^−/−^ mice. GAPDH was used as protein loading control. Densitometric analyses of the immunoblot bands similar to those shown are presented at right (DU: densitometric Unit). Data are shown as means ±SD (*n*=4; *p<0.01 compared to WT). **(b)** p62 immunostained cytospin preparations of sorted CD44^+^Ter119^+^FSC^High^ bone marrow cells from wild-type (WT) and Fyn^−/−^ mice counterstained with DAPI. **Lower Panel.** The puncta mean fluorescence was measured using Image J software. Data are presented as means ±SD (*n*=3); *p<0.05 compared to WT. **(c)** Immunoprecipitates (IP) containing equal amounts of p62 were obtained from sorted CD44^+^Ter119^+^FSC^High^ bone marrow cells from wild-type (WT) and Fyn^−/−^ mice, then subjected to immunoblot with anti-Keap1 or p62 antibody (Wb: Western-blot). The experimental results shown are representative of 4 similar separate experiments. IgG was used as loading control. Densitometric analyses of the immunoblot bands similar to those shown are presented at lower panel (DU: densitometric Unit). Data are shown as means ±SD (*n*=4; *p<0.01 compared to WT). **(d)** Immunoprecipitates (IP) of Keap 1 were obtained from sorted CD44^+^Ter119^+^FSC^high^ bone marrow cells from wild-type (WT) and Fyn^−/−^ mice, then subjected to immunoblot with anti- p62 antibody (Wb: Western-blot). The experimental results shown are representative of 4 similar separate experiments. IgG was used as loading control. Densitometric analyses of the immunoblot bands similar to those shown are presented at lower panel (DU: densitometric Unit). Data are shown as means ±SD (*n*=4; *p<0.01 compared to WT).

**Figure 5.**
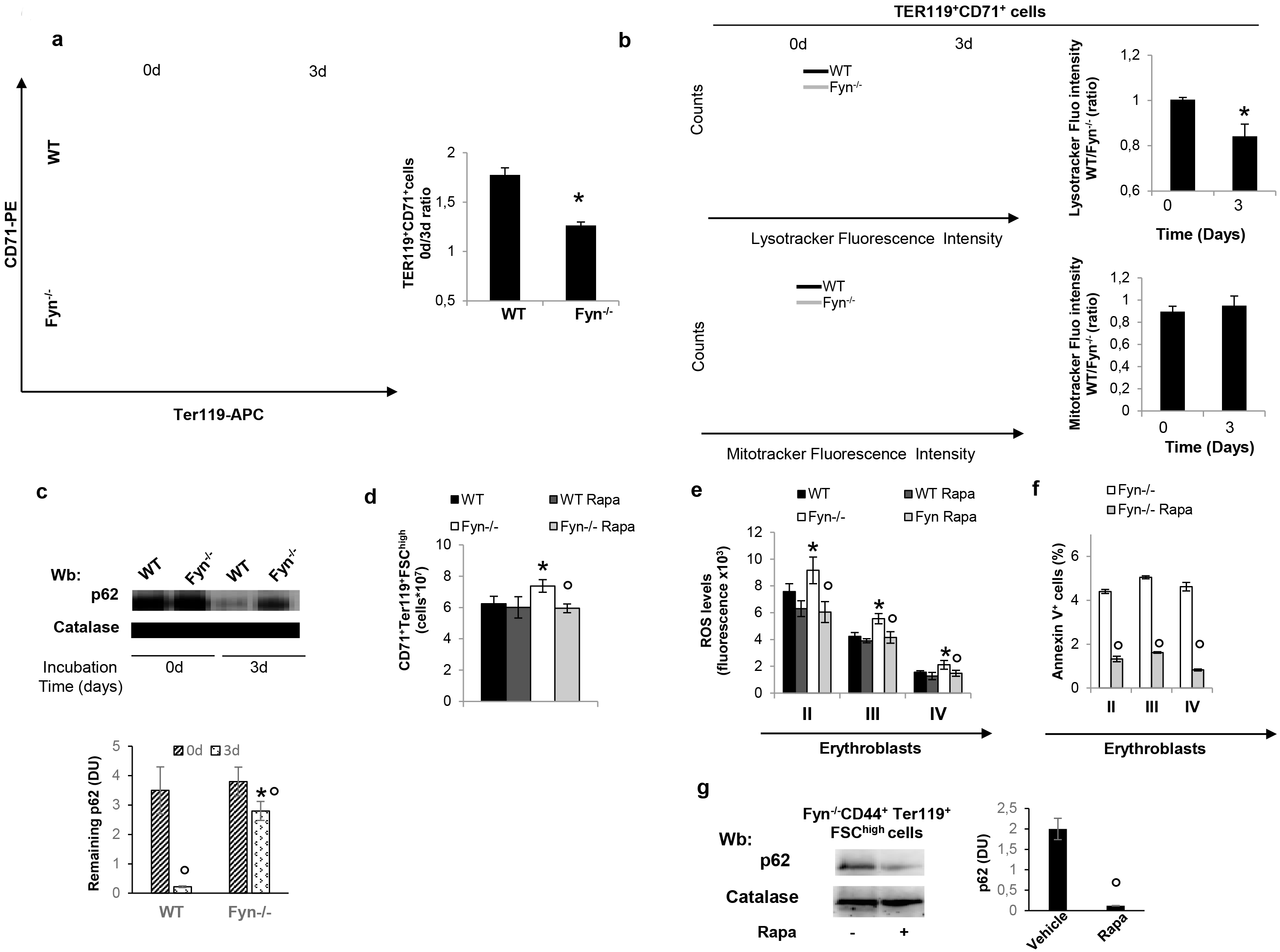
In Fyn^−/−^ mice, the blockage of autophagy results in delayed reticulocyte maturation, decreased lysosomal clearance and accumulation of p62, which is prevented by treatment with Rapamycin, the mTOR inhibitor. **(a)** Flow-cytometric analysis combining CD71, Ter119 and cell size marker strategy on reticulocytes from PHZ treated wild-type (WT) and Fyn^−/−^ mice at baseline (0) and after 3 days of culture. **Right panel.** Bar graph showing the ratio between CD71+/TER119+ at day 0 and 3 in wild-type (WT; *n*=3) and Fyn^−/−^ (*n*=3) mice Data are presented as means ±SD; *p< 0.05 compared to WT mice. **(b)** Flow-cytometry of LysoTracker-stained (upper panel) or MitoTracker-stained (lower panel) at baseline (0) and at 3 days of culture from wild-type (WT) and Fyn^−/−^ mice. One representative of other 3 additional experiments with similar results. **Right panel.** Data are shown as either LysoTracker or MitoTracker fluo intensity WT/Fyn^−/−^ during in vitro reticulocyte maturation (baseline: 0; at 3 days of cell culture). Data are presented as means±SD (*n*=4; *p<0.05 compared to healthy cells). **(c)** Western-blot (Wb) analysis of p62 in maturating reticulocyte at baseline (0) and after 3 days of cell culture. Catalase was used as protein loading control**. Lower panel**. Densitometric analyses of the immunoblot bands similar to those shown are presented at lower panel (DU: densitometric Unit). Data are shown as means ±SD (*n*=3; *p<0.01 compared to WT). **(d)** Effect of treatment with Rapamycin (Rapa) on total erythroid precursors (CD71-Ter119) from the bone marrow of wild-type (WT) and Fyn^−/−^ mice. Data are presented as means ±SD (*n*=6); *p< 0.05 compared to WT; °p< 0.05 compared to vehicle treated animals. **(e-f)** Effect of Rapamycin treatment (Rapa) on ROS levels and Annexin V+ cells in erythroid precursors: population II (Pop II), corresponding to basophilic erythroblasts; population III (Pop III), corresponding to polychromatic erythroblasts and population IV (Pop IV), corresponding to orthochromatic erythroblasts from bone marrow of wild-type (WT) and Fyn^−/−^ mice. Data are presented as means ±SD (*n*=6 from each strain); * p <0.05 compared to WT; °p< 0.05 compared to vehicle treated animals. **(g)** Western-blot (Wb) analysis of p62 in sorted CD44^+^Ter119^+^FSC^high^ bone marrow cells with and without Rapamycin from Fyn^−/−^ mice. Catalase was used as protein loading control. Densitometric analyses of the immunoblot bands similar to those shown are presented at right (DU: densitometric Unit). Data are shown as means ±SD (*n*=4; *p<0.01 compared to WT).

### The mTOR inhibitor Rapamycin unblocks autophagy defect and ameliorates erythropoiesis in Fyn^−/−^ mice

We next tested whether Rapamycin, a known mTOR inhibitor, may modulate Fyn^−/−^ mouse erythropoiesis as reported for other models of pathologic erythropoiesis.^35,36,40,45^ In Fyn^−/−^ mice, Rapamycin administration reduced total erythroblasts, while no significant effects were observed in control animals as previously reported^35,36,40,45^ (Figure 5d). In Fyn^−/−^ mice treated with Rapamycin, this was associated with amelioration of the terminal phase of erythropoiesis (Figure 6Sc). Rapamycin significantly reduced the generation of ROS and the amount of Annexin- V+ positive cells only in erythroid precursors from Fyn^−/−^ mice (Figure 5e, 5f). In agreement with Rapamycin induced activation of autophagy, we found a significant reduction in levels of p62 in erythroblasts from Rapamycin treated Fyn^−/−^ mice compared to vehicle treated animals (Figure 5g).

We also evaluated the impact of anti-oxidant treatment with N-acetylcysteine (NAC), which has been shown to indirectly modulate autophagy by reducing intracellular oxidation.^46,47^ In Fyn^−/−^ mice, NAC reduced total number of erythroblasts, increased orthochromatic erythroblasts and reduced the amount of Annexin V+ cells, indicating an amelioration of erythropoiesis in Fyn^−/−^ mice (Figure 7Sa, b, c). The findings indicate that Rapamycin unblocks autophagy, allowing the degradation of accumulated proteins and ameliorating erythropoiesis in Fyn^−/−^ mice.

### Rapamycin rescues the abnormal response of Fyn^−/−^ erythroblasts to stress erythropoiesis

Next, we investigated whether Rapamycin may alleviate the PHZ-induced stress erythropoiesis in Fyn^−/−^ mice. As shown in Figure 6a, the administration of Rapamycin in PHZ treated Fyn^−/−^ mice resulted in milder anemia and faster recovery compared to PHZ treated Fyn^−/−^ mice. Since at day 4 following PHZ administration a plateau in Hct value was evident in both Fyn^−/−^ mouse groups, we focused on this time point to carry out the analysis of erythropoiesis in both mouse strains exposed to either PHZ or PHZ and Rapamycin. As shown in Figure 6b, we found decreased extramedullary splenic erythropoiesis but increased bone marrow erythropoiesis in PHZ-Rapamycin treated Fyn^−/−^ mice compared to PHZ treated animals.

**Figure 6.**
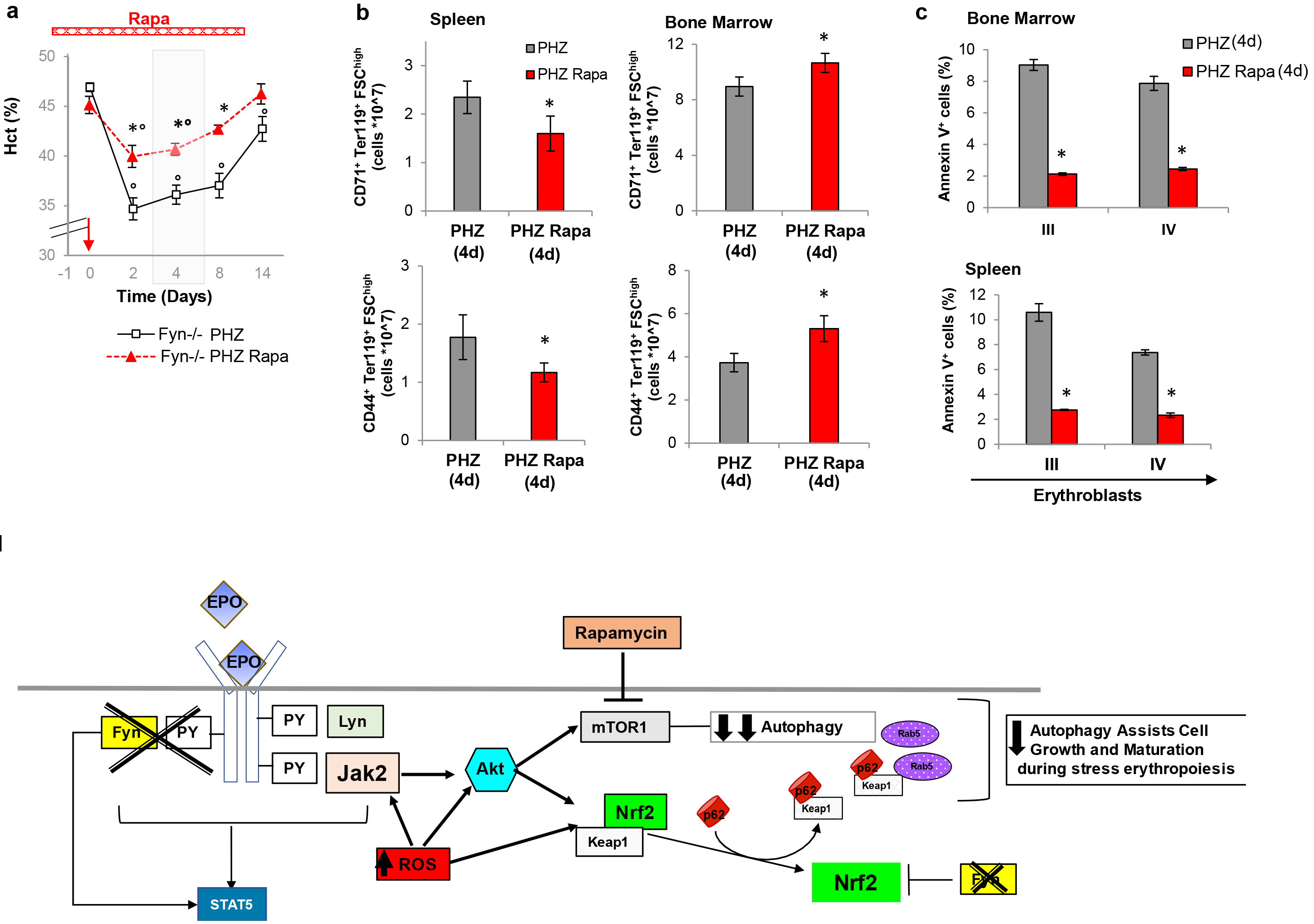
**(a)** Hematocrit (%) in Fyn^−/−^ (*n*=6) mice treated with either phenylhydrazine alone (PHZ) or combined with Rapamycin (Rapa- red bar is the time of treatment with it). The red arrow indicates the injection of PHZ. Data are presented as means ±SD; *p< 0.05 compared to PHZ treated animals; °p<0.05 compared to baseline values. The gray area identifies the window time for characterization of the stress erythropoiesis in Fyn^−/−^ mice. **(b)** Cyto-fluorimetric analysis of total erythroid precursors from either bone marrow or spleen of Fyn^−/−^ mice treated with either PHZ alone or Rapamycin (Rapa) plus PHZ (4 days after injection). Results from CD44-Ter119 (lower panel) or CD71Ter119 (upper panel) strategies are shown. Data are presented as means ±SD (*n*=4); *p< 0.05 compared to PHZ treated animals; °p<0.05 compared to baseline values. **(c)** Amount of Annexin V^+^ cells in population III (Pop III), corresponding to polychromatic erythroblasts and population IV (Pop IV), corresponding to orthochromatic erythroblasts from either spleen or bone marrow of Fyn^−/−^ mice. Data are presented as means ±SD (*n*=4 from each strain); *p<0.05 compared to WT. **(d)** Schematic diagram of the role of Fyn in stress erythropoiesis. Our data show that Fyn is part of the EPO signaling cascade, directly affecting EPO-receptor (R) Tyr-phosphorylation as well as the activation of STAT5. Thus, Fyn acts also as downstream regulator of EPO signaling pathway. The increase oxidation related to reduced efficiency of EPO signaling promotes oxidation, which activates Jak2-Akt-mTOR pathway, resulting in suppression of autophagy. This is required to assist cell growth and maturation during both normal and in particular stress erythropoiesis. The dysregulation of Nrf2 due to the absence of Fyn, which is involved in Nrf2 post-induction repression, promote the accumulation of proteins such as the Keap1-p62 complex, further supporting the blockage of autophagy in erythroblasts genetically lacking Fyn.

The amount of Annexin-V^+^ cells was also markedly reduced by the co-administration of Rapamycin and PHZ in Fyn^−/−^ mouse erythroblasts from both bone marrow and spleen compared to PHZ alone (Figure 6c). It is interesting to note that, in wild-type animals the co-administration of Rapamycin worsened the PHZ induced stress erythropoiesis as previously reported^36^ (Figure 8Sa, b, c). Collectively, these data support the idea that by activating autophagy it is possible to rescue the altered response of Fyn^−/−^ mice to stress erythropoiesis.

## DISCUSSION

We identified Fyn as a new kinase involved in EPO signaling cascade during normal and stress erythropoiesis following treatments with PHZ and Doxorubicin to induce acute anemia. Previous studies have documented a similar role for Jak2 and Lyn kinases in erythropoiesis.^2–4,48,49^ The reduction in EPO induced phosphorylation of STAT5 noted in Fyn^−/−^ mouse erythroid cells implies a role for Fyn downstream of STAT5 and suggests that Fyn is implicated in EPO signaling pathway by intersecting the activation of EPO-R through STAT5. Similar findings have been reported for mice genetically lacking Lyn,^48^ supporting the important and non-redundant role of Fyn in EPO mediated signaling cascade. The also suggest that multiple kinases may be important in coordinating stress erythropoiesis in health and disease.

In Fyn^−/−^ mice, the reduced effectiveness of EPO signaling results in increase ROS generation and cell apoptosis. The attempt of Fyn^−/−^ mouse erythroblasts to adapt to oxidative stress is indicated by the activation of the redox related transcription factor Nrf2.^33,35^ However, the persistent activation of Nrf2 due to the absence of its physiologic repressor Fyn, resulted in accumulation of damaged/non-functional proteins that further amplified the intracellular oxidative stress.^13,14^ Indeed, Fyn ^−/−^ mice display up-regulation of several cytoprotective systems, such as HO-1, but without effective functioning of these cytoprotective systems, indicating an impairment of the protein quality control process mediated by autophagy.

Previous studies have shown that perturbation of autophagy is detrimental for erythroid maturation and has been documented in condition with disordered erythropoiesis such as β-thalassemic syndromes or chorea-acanthocytosis or during iron deficiency.^18,35,44,50,51^ In Fyn ^−/−^ mice, the ROS mediated activation of Jak2-Akt-mTOR pathway represses autophagy and thereby contributes to the ineffective erythropoiesis of Fyn ^−/−^ mice.^4,37^ Consistent with this hypothesis, in Fyn ^−/−^ mouse erythroblasts we found accumulation of the autophagy cargo protein p62, a marker of autophagy inhibition. In addition, p62 was also complexed with Keap1, the Nrf2 shuttle protein,^14^ further supporting the dysregulation of Nrf2 and the blockage of autophagy during Fyn^−/−^ erythropoiesis. The accumulation of Rab5, a small GTPase involved in endocytic vesicular transport,^52^ suggested a possible perturbation of the autophagy lysosomal system in Fyn^−/−^ mice. Indeed, we found a reduction in lysosome clearance during Fyn^−/−^ mouse reticulocyte maturation in presence of preserved clearance of mitochondria compartment.

The ability of Rapamycin, a known mTOR inhibitor and autophagy activator, to ameliorate Fyn^−/−^ mouse baseline erythropoiesis and to prevent accumulation of p62 support the blockage of autophagy in Fyn ^−/−^ mice. This is further corroborated by the observation that Rapamycin co-administrated with PHZ restored the erythropoietic response in Fyn^−/−^ mice. These finding shed a new light on the link between dysregulation of Nrf2 and impairment of autophagy in stress erythropoiesis, demonstrating the multimodal action of Fyn in establishing the developmental program of erythropoiesis.

In conclusion, our data indicate Fyn as a new kinase important in erythropoiesis, contributing to activation of EPO cascade, further increasing the complexity of EPO signaling pathway. The absence of Fyn generates a “domino effect” of the cell backup mechanisms in response to increase oxidative stress induced by the reduced efficiency of EPO signal. The dysregulation of post-induction repression Nrf2 results in accumulation of aggregated proteins, which further increase cellular oxidative stress. This promotes the activation of Jak2-Akt-mTOR pathway with repression of autophagy, which is required to assist cell growth and maturation in normal and stress erythropoiesis. The rescue experiments with Rapamycin co-administrated to PHZ further reinforce the importance of autophagy as adaptive mechanism to stressful condition in presence of perturbation of EPO signaling pathway. Future studies will be required to fully characterize the signaling pathways intersected by Fyn during pathologic erythropoiesis.

## METHODS

### Mouse strains and design of the study

The Institutional Animal Experimental Committee of University of Verona (CIRSAL) and the Italian Ministry of Health approved the experimental protocols. Two-months old female wild-type (WT) and Fyn^−/−^ mice were studied. Where indicated, WT and Fyn^−/−^ mice were treated with EPO (10 U/mouse/day for 5 days by intraperitoneal injection),^3^ or Phenylhydrazine (PHZ: 40 mg/Kg on day 0 by intraperitoneal injection),^17^ or Doxorubicin (DOXO: 0.25 mg/Kg on day 0 by intraperitoneal injection)^19^ to study stress erythropoiesis. Rapamycin (Rapa) was administrated at the dosage of 10 mg/Kg/d by intraperitoneal injection for 1 week, then mice were analyzed. In experiments with PHZ co-administration, Rapa was given at the dosage of 10 mg/Kg/d by intraperitoneal injection one day before PHZ administration (40 mg/Kg body; single dose at day 0) and then Rapa was maintained for additional 14 days. N-Acetylcysteine (NAC, 100 mg/Kg body; intraperitoneally injected) was administrated for 3 weeks as antioxidant treatment.^16,17^ In mouse strains, hematological parameters, red cell indices and reticulocyte count were evaluated at baseline and at different time points (6, 8 and 11 days after EPO injection; at 2, 4, 8 and 14 days after PHZ injection; at 3, 6 and 9 days after DOXO injection; at 2, 4, 8, 14 days after Rapa plus PHZ injection) as previously reported.^20,21^ Blood was collected with retro-orbital venipuncture in anesthetized mice using heparinized microcapillary tubes. Hematological parameters were evaluated on a Bayer Technicon Analyser ADVIA. Hematocrit and hemoglobin were manually determined.^22,23^

### Flow cytometric analysis of mouse erythroid precursors and molecular analysis of sorted erythroid cells

Flow cytometric analysis of erythroid precursors from bone marrow and spleen from WT and Fyn^−/−^ was carried out as previously described using the CD44-Ter119 or CD71-Ter119 strategies.^16,24,25^ Analysis of apoptotic basophilic, polychromatic and orthochromatic erythroblasts was carried out on the CD44-Ter119 gated cells using the Annexin-V PE Apoptosis detection kit (eBioscience, San Diego, CA, USA) following the manufacturer’s instructions. Erythroblasts ROS levels were measured as previously reported by Matte et al.^16^ Sorted cells were used for (i) morphological analysis of erythroid precursors on cytospin preparations stained with May Grunwald-Giemsa; (ii) immuno-blot analysis with specific antibodies against anti-P-Ser473-Akt, anti-Akt, anti-P-Ser2448-mTOR, anti-mTOR, anti-Jak2 (Cell Signaling, Massachusetts, USA); anti-P-Ser40-Nrf2, anti-Nrf2, anti-p62, anti-Rab5 (Abcam, Cambridge, UK); anti-Keap1 (Proteintech, Manchester, UK); anti-EPO-R (Sigma-Aldrich, Missouri, USA); anti-STAT5, anti-Lyn (Santa Cruz Biotechnology, Texas, USA); anti-GAPDH (Santa Cruz Biotechnology, Texas, USA) and anti-catalase (Abcam, Cambridge, UK) were used as loading control; (iii) immunoprecipitation assay; and (iv) RT-PCR analysis. Details of immunoprecipitation, RT-PCR and immuno-blot protocols used for the analysis of sorted erythroblasts are described in Supplementary materials and methods.

### CFU-E, BFU-E assay

CFU-E and BFU-E assay was carried out using MethoCult as previously reported.^26^ Details are present in Supplementary Methods.

### Immunofluorescence assay for p62 and FOXO3 in sorted erythroblasts

Immunofluorescence assay for p62 and FOXO3 in sorted erythroblasts was carried out as previously described.^18,23,27^ Details are reported in Supplementary materials and methods.

### LysoTracker and MitoTracker analysis in maturating reticulocytes

To obtain reticulocyte enriched RBC fraction, WT and Fyn^−/−^ mice were intraperitoneally injected with PHZ (40 mg/kg) at day 0, 1, 3 to induce reticulocytosis, and blood was collected in heparinized tubes at day 7, as previously described.^28^ RBCs were washed three times with the maturation medium (60% IMDM, 2mM L-glutamine, 100U Penicillin-Streptomicin, 30% FBS, 1% BSA and 0.5 μg/ml Amphotericin), diluted 1/500 in maturation medium and cultured in a cell culture incubator at 37°C, 5% of CO_2_ for 3 days. Clearance of Lysosome and Mitochondria, on the CD71/Ter119 gated RBC populations, were analyzed at day 0 and 3 of culture using the Lysotracker Green DND-26 (ThermoFisher Scientific) and the MitoTracker Deep Red (ThermoFisher Scientific) probes, respectively, following the manufacturer’s instructions. Samples were acquired using the FACSCantoll™ flow cytometer (Becton Dickinson, San Jose, CA, USA) and data were processed with the FlowJo software (Tree Star, Ashland, OR, USA) as previously described.^16,17^

### Pearl’s analysis of liver and spleen

Immediately following dissection, spleen and liver were formalin-fixed and paraffin-embedded for Pearl’s staining.

### Molecular analysis of liver

Protocols used for RNA isolation, cDNA preparation and quantitative qRT-PCR have been previously described.^29^ Detailed primer sequences are available on request and shown in Table 1S. Liver immuno-blot analysis was performed as previously described.^16,30^

### Measurement of heme and heme-oxygenase-1 activity

Liver heme content was measured using a fluorescence assay, as previously reported.^31^ Details are reported in Supplementary Methods.

HO activity was evaluated in tissue microsomal fractions by spectrophotometric determination of bilirubin produced from hemin added as the substrate, as previously reported.^32^

### Statistical analysis

Data were analyzed using either t-test or the 2-way analysis of variance (ANOVA) for repeated measures between the mice of various genotypes. A difference with a p< 0.05 was considered significant.

## ACKNOWLEDGMENTS

This work was supported by FUR-UNIVR (LDF).

## AUTHORSHIP AND CONTRIBUTIONS

LDF, AI designed the experiments, analyzed the data and wrote the manuscript; ET, SG, NM contributed to study design and the writing of the manuscript; EB, AM contributed in designing the experiments, carried out the experiments, analyzed the data and contributed in writing the manuscript; SCMT carried out CFU-E-BFU-E assay; DC, SP performed the experiments on heme; VM carried out the immunomicroscopy on FOXO3; LD carried out the molecular analysis and contributed to data analysis. AS, EF and ABW contributed to immunoblot experiments.

## THE PAPER EXPLAINED

### Problem

Erythropoiesis is a complex multistep process responsible of the production of circulating mature erythrocytes and involved the production of reactive oxygen species (ROS) during erythroid differentiation. Although progresses have been made in identification of molecular mechanisms involved in normal erythropoiesis, much still remains to be investigated in cellular response to stress erythropoiesis. Word-wide distributed hereditary red cell disorders such as β-thalassemic syndromes are characterized by ineffective erythropoiesis, known model of stress erythropoiesis. Growing evidences involve kinases of the Src family (SFK) in receptor signaling pathways in cell growth and maturation of hematopoietic cells. While several studies have shown Fyn to be involved in megakariocytosis and STAT5 regulation, the role of Fyn in normal and stress erythropoiesis is still unknown.

### Results

Here, we show that the absence of Fyn promotes diserythropoiesis and increased oxidation of erythroblasts from Fyn^−/−^ mice, related to a reduced efficiency of erythropoietin (EPO) signaling cascade. We demonstrated that Fyn targets EPO-receptor and is involved in modulation of STAT5, as additional kinase to the canonical Jak2 and Lyn kinase. Using phenylhydrazine (PHZ) or doxorubicine (Doxo) to induce stress erythropoiesis, we found a blunted recovery and prolonged anemia in Fyn^−/−^ mice, supporting Fyn as downstream effecteor on EPO cascade. In Fyn^−/−^ mice, the increase ROS levels promotes Jak2-Akt overactivation targeting the redox-sensitive transcriptional factor Nrf2 and mTOR, the gatekeeper of autophagy. Since Fyn is involved in post-induction regulation of Nrf2, we show persistent activation of Nrf2 and accumulation of non-functional proteins. This was also favored by mTOR mediated suppression of autophagy, leading to perturbation of reticulocyte maturation and lysosomal clearance.

We demonstrate that Rapamycin, targeting mTOR, improved autophagy and reduced apoptosis, restoring Fyn^−/−^ mouse erythropoietic response to PHZ stress.

### Impact

Our findings define a novel role of Fyn in EPO signaling cascade and underscore the importance of Fyn in modulating cellular response to oxidation. Since we show the multimodal action of Fyn in the developmental program of erythropoiesis, our data point out Fyn as potential new leading factor in diseased erythropoiesis linked to oxidative stress.

